# Inhibitory concentrations of ciprofloxacin induce an adaptive response promoting the intracellular survival of *Salmonella* Typhimurium

**DOI:** 10.1101/2021.05.06.443048

**Authors:** Sushmita Sridhar, Sally Forrest, Derek Pickard, Claire Cormie, Emily Lees, Nicholas R Thomson, Gordon Dougan, Stephen Baker

## Abstract

Antimicrobial resistance (AMR) is a pressing global health crisis, which has been fuelled by the sustained use of certain classes of antimicrobials, including fluoroquinolones. While the genetic mutations responsible for decreased fluoroquinolone (ciprofloxacin) susceptibility are known, the implications of ciprofloxacin exposure on bacterial growth, survival, and interactions with host cells are not well described. Aiming to understand the influence of inhibitory concentrations of ciprofloxacin *in vitro*, we subjected three clinical isolates of *S.* Typhimurium to differing concentrations of ciprofloxacin, dependent on their minimum inhibitory concentrations (MIC), and assessed the impact on bacterial growth, morphology, and transcription. We further investigated the differential morphology and transcription that occurred following ciprofloxacin exposure and measured the ability of ciprofloxacin-treated bacteria to invade and replicate in host cells. We found that ciprofloxacin-exposed *S.* Typhimurium are able to recover from inhibitory concentrations of ciprofloxacin, and that the drug induces specific morphological and transcriptional signatures associated with the bacterial SOS response, DNA repair, and intracellular survival. In addition, ciprofloxacin-treated *S.* Typhimurium have increased capacity for intracellular replication in comparison to untreated organisms. These data suggest that *S.* Typhimurium undergoes an adaptive response under ciprofloxacin perturbation that promotes cellular survival, a consequence that may justify more measured use of ciprofloxacin for *Salmonella* infections. The combination of multiple experimental approaches provides new insights into the collateral effects that ciprofloxacin and other antimicrobials have on invasive bacterial pathogens.

**Importance:** Antimicrobial resistance is a critical concern in global health. In particular, there is rising resistance to fluoroquinolones, such as ciprofloxacin, a first-line antimicrobial for many Gram-negative pathogens. We investigated the adaptive response of clinical isolates of *Salmonella* Typhimurium to ciprofloxacin, finding that the bacteria adapt in short timespans to high concentrations of ciprofloxacin in a way that promotes intracellular survival during early infection. Importantly, by studying three clinically relevant isolates, we were able to show that individual isolates respond differently to ciprofloxacin, and for each isolate, there was a heterogeneous response under ciprofloxacin treatment. The heterogeneity that arises from ciprofloxacin exposure may drive survival and proliferation of *Salmonella* during treatment and lead to drug resistance.

## Introduction

The current trajectory of resistance to numerous broad-spectrum antimicrobials in bacterial pathogens is steadily increasing, making antimicrobial resistance (AMR) of critical concern for human health. This problem is further exacerbated by the fact there are few novel antimicrobials in the developmental pipeline and a dearth of vaccines to prevent against the increasing number of drug-resistant bacterial infections (1, 2). A large burden of multi-drug resistant (MDR) organisms arise in low-middle income countries (LMICs), which, in part, may be associated with high level usage of broad-spectrum antimicrobials in the community (1). These factors pose a serious global health threat.

Fluoroquinolones are amongst the most commonly used broad-spectrum antimicrobials globally, and are commonly administered for urinary tract infections, pneumonia, dysentery, and febrile diseases (3). This potent group of bactericidal chemicals act by binding to bacterial type II topoisomerases DNA gyrase I (GyrA and GyrB) and Topoisomerase IV (ParC and ParE) to disrupt DNA supercoiling, which leads to cell death (4–6). Resistance to fluoroquinolones is associated with specific mutations in the *gyrA, gyrB, parC* and/or *parE* genes, although the extent of resistance can be further modulated by mutations in efflux pumps and porins and also via the acquisition of plasmid-mediated quinolone resistance (PMQR) genes (7–11). Ciprofloxacin is the most widely available fluoroquinolone, and its common use, particularly in LMICs, has resulted in widespread resistance in once-susceptible pathogens (12–14).

Despite extensive resistance, ciprofloxacin remains commonly used, and given its mode of action, it is likely to induce a range of additional cellular responses (15). Transcriptional studies of bacteria exposed to ciprofloxacin have shown that pathways associated with the stress response, solute and drug transport, DNA repair, and phage induction are upregulated, which can increase error-prone DNA replication and bacterial resilience during ciprofloxacin exposure (16–21). However, little is known about how bacterial genotype influences the response to ciprofloxacin or how bacteria respond when exposed to inhibitory concentrations of the drug.

*Salmonella* enterica serovar Typhimurium (*S.* Typhimurium) is a Gram-negative enteric bacterium that typically causes a self-limiting gastroenteritis in humans but is also associated with invasive disease in the immunocompromised. Specific *S.* Typhimurium lineages, such as ST313 and ST34, are associated with invasive disease in parts of sub-Saharan Africa and Southeast Asia, respectively, and have independently developed resistance to ciprofloxacin multiple times (22, 23). AMR in these organisms typically arises during local outbreaks and is not ubiquitous, demonstrating that there can be variation in the AMR profile of individual bacteria belonging to a single clade.

Although the cellular mechanisms of ciprofloxacin resistance are well-defined, our understanding of how bacteria evolve and adapt during short-term exposure to ciprofloxacin is limited. Specifically, there is a lack of evidence regarding how clinical isolates respond to antimicrobials they may commonly encounter indirectly during therapy. Here, aiming to understand how genotypic and phenotypic characteristics are impacted by fluoroquinolone exposure, we studied three diverse *S.* Typhimurium variants (an ST313, ST34, and ST19), under sustained perturbation with ciprofloxacin. Focusing on an ST313 isolate, we found that bacteria have substantial resilience to high concentrations of ciprofloxacin and that ciprofloxacin-exposed bacteria undergo distinct morphological and transcriptional changes within a short timeframe, impacting on bacterial survival and their interactions with host cells.

## Results

### Salmonella Typhimurium can replicate in inhibitory concentrations of ciprofloxacin

Using three isolates of *S.* Typhimurium selected by sequence type (ST) and ciprofloxacin susceptibility, time kill curves were performed in the presence of 0×, 1×, 2×, and 4× the ciprofloxacin MIC of each isolate to determine growth dynamics over a 24-hour period of ciprofloxacin exposure (**Figure 1**). Quantification of colony forming units (CFU) demonstrated that bacterial growth was most likely to be inhibited between 0-and 6-hours post-exposure, and the rate of growth was dependent on ciprofloxacin concentration. However, after six hours of ciprofloxacin exposure, there was a “recovery” phase, during which bacteria in the treated conditions began to replicate and increase in CFU (**Figure 1A–C**). This trend was observed in all three isolates, although the degree of recovery and absolute number of organisms between isolates was variable. *S.* Typhimurium D23580 (ST313) bacteria showed the largest range of growth responses to different ciprofloxacin concentrations (**Figure 1A**). All conditions of SL1344 (ST19) and VNS20081 (ST34) bacteria had comparable CFUs at 24 hours, whereas there was considerably more variation in D23580 by treatment and replicate (**Figure 1D–F**). In addition, after eight hours exposure at 2× MIC of ciprofloxacin, the mean cellular concentration of D23580 was 153 ± 217 CFU/ml; in analogous conditions at 24 hours, the mean CFU/ml was 46,000 ± 65,000 (**Figure 1A, D**). The lower variability in SL1344 and VNS20081 cultures may be explained by their genetic backgrounds or specific ciprofloxacin MIC. Notwithstanding the experimental variation observed in D23580 cultures, the overall trend across multiple replicates of the three isolates was that bacteria under high ciprofloxacin exposure were able to reach a concentration comparable to non-treated bacteria after 24 hours.

**Figure 1.**
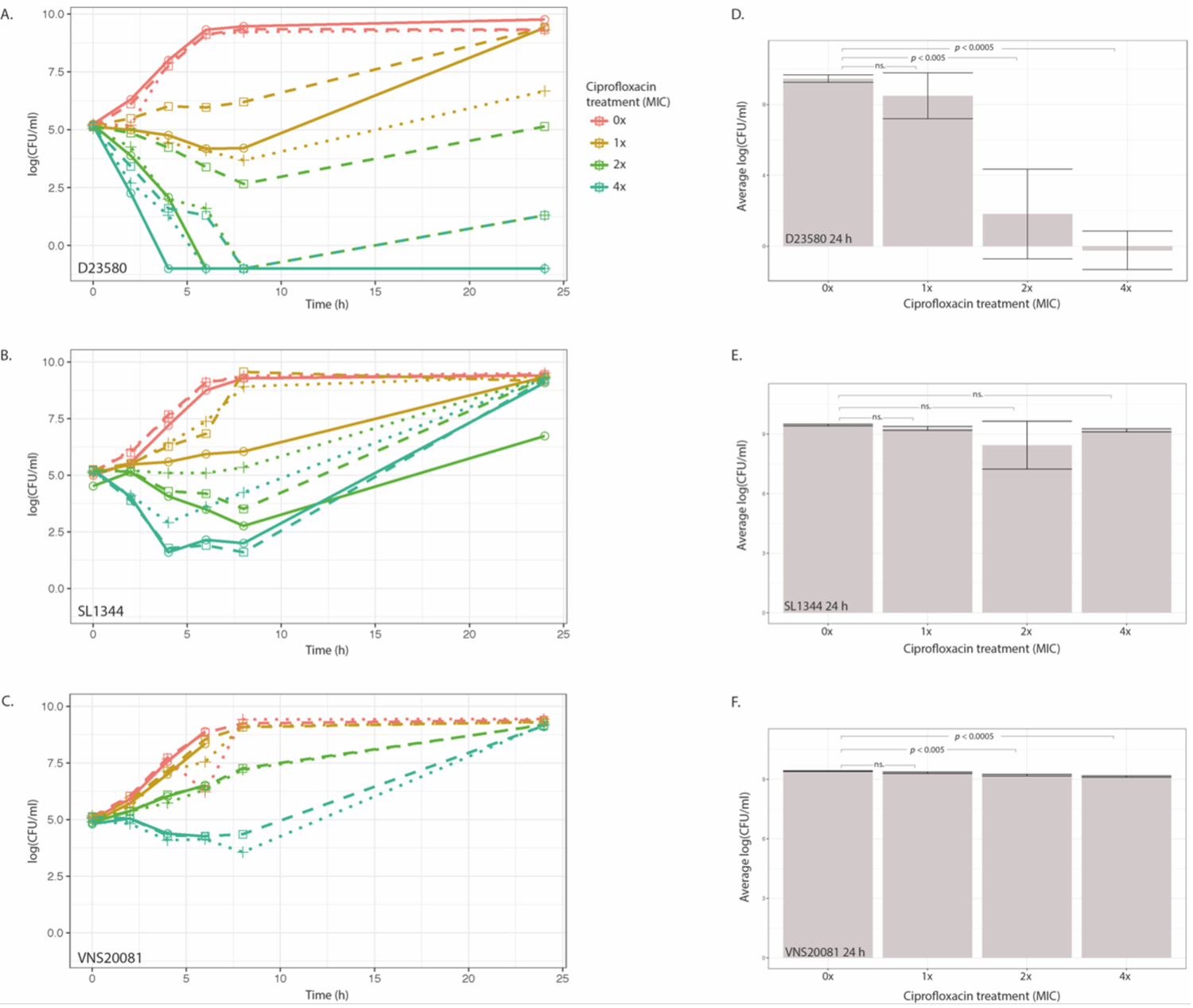
Time kill curves of *S.* Typhimurium isolates at different ciprofloxacin concentrations. *S.* Typhimurium isolates D23580 (**A**), SL1344 (**B**), and VNS20081 (**C**) were grown for 24 hours in four concentrations of ciprofloxacin (0×, 1×, 2×, or 4× MIC) and subjected to CFU enumeration at 6 time points post-inoculation. Three biological replicates were performed, and each replicate was plotted independently. The average CFU/ml was calculated for each isolate and condition for the 24-hour time point and plotted as mean ± SD (**D**, D23580; **E**, SL1344; **F**, VNS20081). An ANOVA was performed to compare means at 24 hours, and Dunnett’s test was performed to compare 24 hour means of 1×, 2×, and 4× ciprofloxacin MIC to 0× (control).

We postulated that this recovery in growth was associated with ciprofloxacin degradation. To assess this, we centrifuged and filter-sterilised the ciprofloxacin-containing media after 24 hours of bacterial growth. D23580 was inoculated into this filter sterilised media and incubated at 37°C for a further 24 hours, as before. The time kill curves replicated those of the original assays, indicating that the inhibitory activity of ciprofloxacin was preserved at approximately the same concentrations for the same time periods (**Figure S1**).

To determine whether the recovery of organisms under ciprofloxacin treatment was due to acquired mutations, we performed whole genome sequencing on D23580 grown for 24 hours without antimicrobial supplementation or with 0.06 μg/ml ciprofloxacin (2× MIC) to detect dominant SNPs. To capture the genetic signatures of culturable organisms only, bacteria were grown in liquid broth for 24 hours and then spread on agar plates. Colonies were pooled from each plate for DNA extraction and sequencing. Aiming to identify dominant mutations arising in D23580 across three biological replicates, we found mutations in *ramR* and *gyrA* in bacteria grown in ciprofloxacin. There were no mutations in untreated D23580. The occurrence of SNPs in *ramR* suggests that this gene plays a critical role in modulating bacterial survival during exposure to high ciprofloxacin concentrations in the absence of *gyrA* mutations (**Table 1**). Notably, only one of the three ciprofloxacin-treated cultures gained a *gyrA* mutation, which highlights the importance of studying other factors that may contribute to bacterial survival upon exposure to high doses of ciprofloxacin.

**Table 1.**
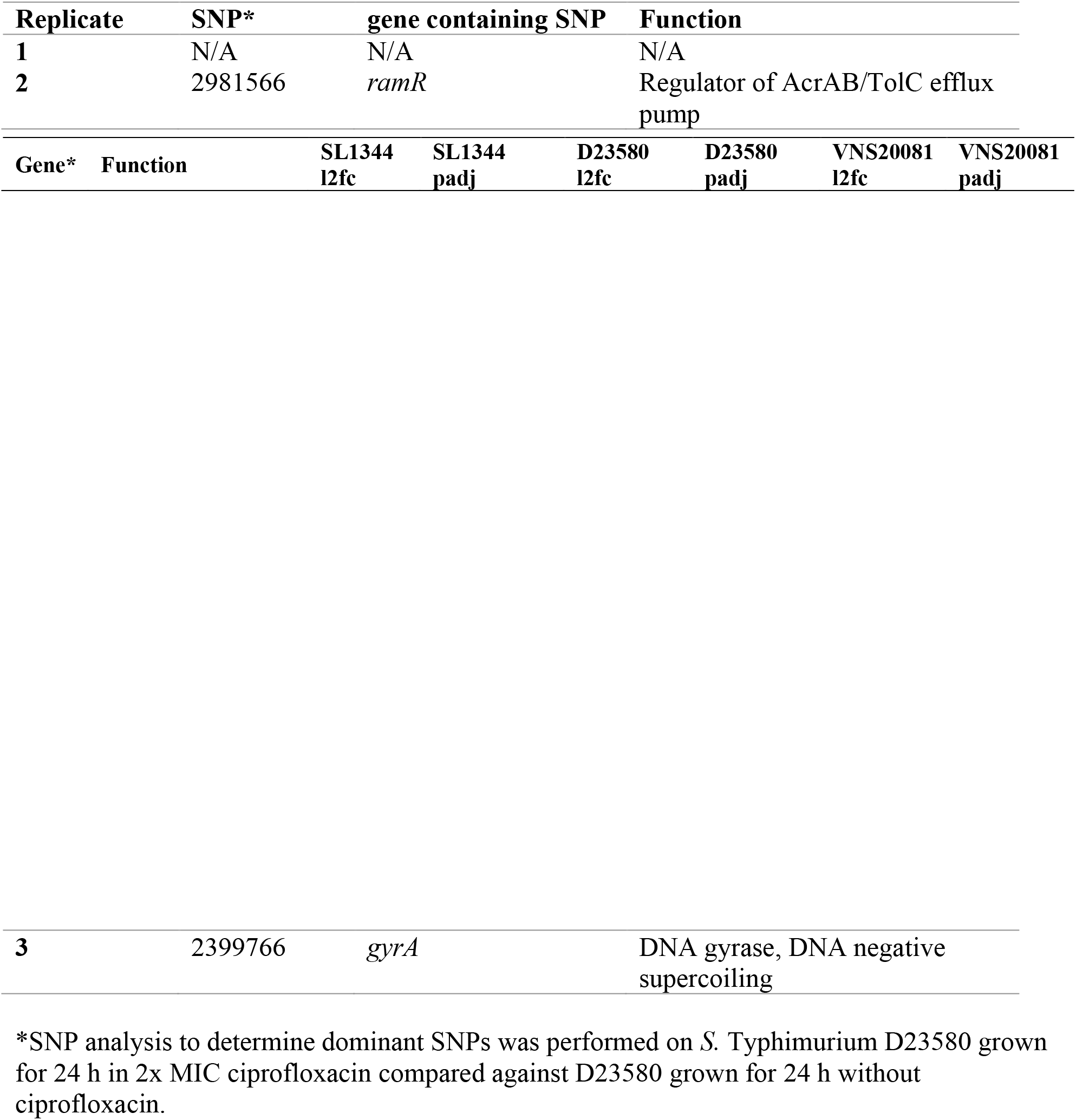
Dominant SNPs found after 24 h growth in 2× MIC ciprofloxacin.

### Ciprofloxacin induces morphological changes in S. Typhimurium

To better understand the impact of ciprofloxacin on the selected organisms, we exposed organisms D23580, SL1344, and VNS20081 to 0×, 1×, 2×, or 4× MIC of ciprofloxacin for two hours and then imaged them using a quantitative high content confocal microscopy system (24) (S. Sridhar and S. Forrest, submitted for publication). A timepoint of two hours was selected to capture early adaptive response. We found that the majority of ciprofloxacin-treated bacteria developed an elongated morphology within two hours of ciprofloxacin exposure (**Figure 2**). Some diversity in bacterial length upon ciprofloxacin treatment was apparent, suggesting a heterogeneous response to ciprofloxacin exposure. Quantitative image analysis of the lengths of individual organisms after two hours indicated substantial heterogeneity in ciprofloxacin-exposed organisms and untreated bacteria (**Figure 2B–D**). However, the mean length of non-treated bacteria was significantly less than that of ciprofloxacin-treated bacteria (D23580: 3.24 μm (0×), 6.73 (1×), 6.40 (2×), 6.13 (4×), *p* < 0.001; SL1344: 2.89 (0×), 4.82 (1×), 6.73 (2×), 6.70 (4×), *p* < 0.001; VNS20081: 3.24 μm (0×), 4.92 (1×), 6.54 (2×), 7.23 (4×), *p* < 0.001). Additionally, there also appeared to be variation in mean and maximum lengths of bacteria between the three isolates, with 4× MIC VNS20081 showing the greatest quantifiable change from untreated VNS20081 (mean of 7.23 μm versus 3.24 μm) (**Figure 2D**). Moreover, there was not a uniform density distribution of cellular lengths; this observation was particularly apparent in the ciprofloxacin-treated bacteria. In particular, a number of 2× MIC ciprofloxacin-treated D23580 and VNS20081 bacteria elaborated considerable elongation, with lengths >30 μm (**Figure 2 B, D**). Such a wide distribution of bacterial lengths indicates that ciprofloxacin exposure drives the formation of discrete bacterial populations of variable lengths.

**Figure 2.**
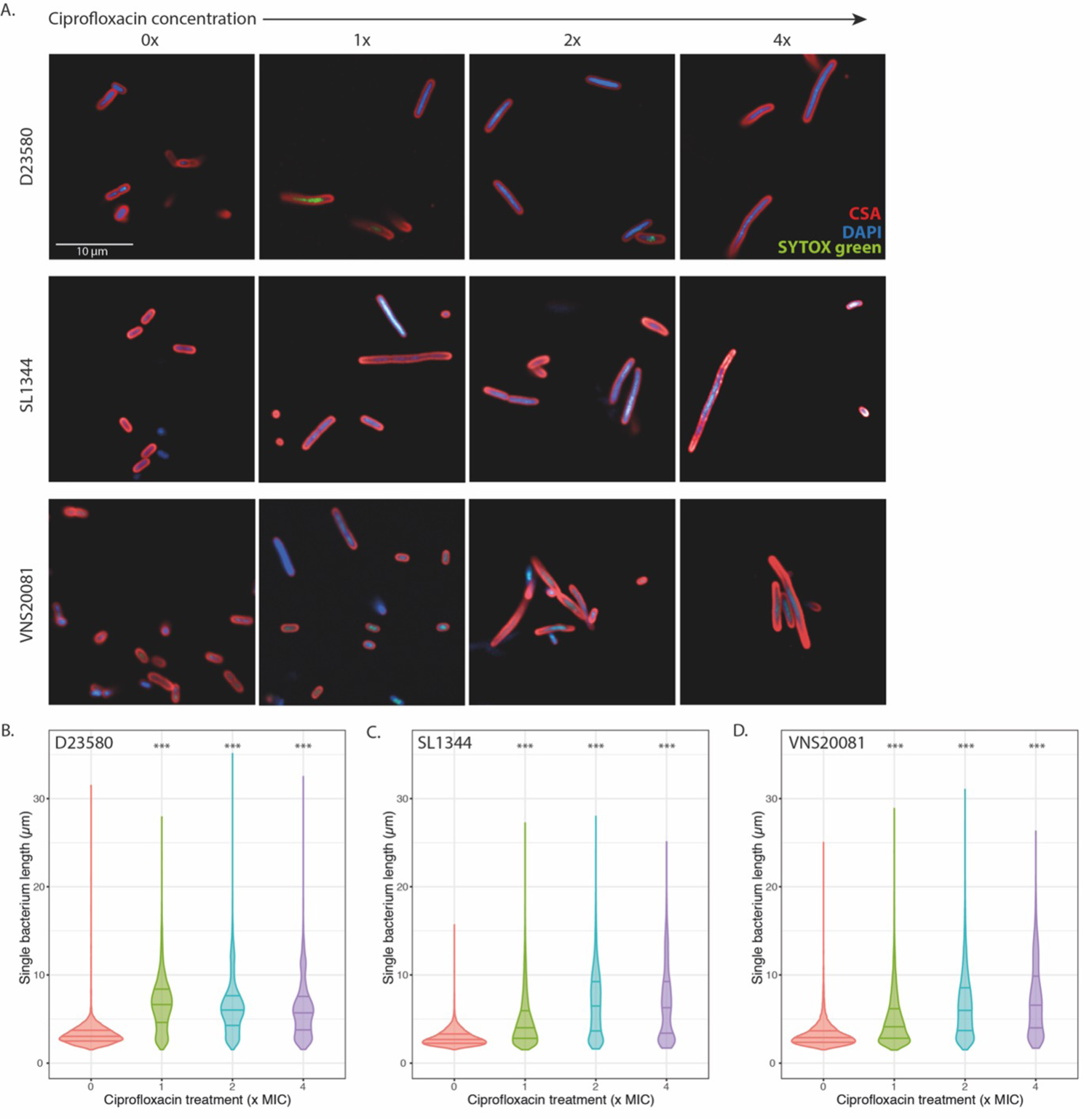
Imaging of *S.* Typhimurium following 2 hours of ciprofloxacin exposure. **A.** D23580 (top panel), SL1344 (middle panel), and VNS20081 (bottom panel) were subjected to 4 concentrations of ciprofloxacin (0×, 1×, 2×, or 4× MIC) and stained and imaged using an Opera Phenix high content microscope. Bacterial membranes were stained using CSA (red), nucleic acids were stained using DAPI (blue), and permeabilized, dead cells were stained using SYTOX green (green). Imaging experiments were carried out in triplicate, with two technical replicates; images from one replicate shown. **B.** The length of single bacteria (μm) was measured quantitatively based on image analysis, and these were plotted for each isolate and condition independently (D23580, left panel; SL1344, middle panel; VNS20081, right panel). Bacterial lengths were plotted as median and interquartile ranges, and the mean ± SD was calculated for each condition compared to 0x-MIC treatment. One-way ANOVAs were performed. * *p* < 0.05. 0x-treated bacteria are in red; 1x-treated bacteria are in green; 2x-treated bacteria are in blue; and 4x-treated bacteria are in purple.

### Ciprofloxacin triggers isolate-specific transcriptional responses

Aiming to investigate the transcriptional features in the chromosome that may induce changes in survival and morphology, total RNA was extracted from the three isolates after two hours exposure to 2× MIC of ciprofloxacin and subjected to sequencing. This time point was selected to best capture early responses before significant cell death. Generally, the broad transcriptional profile was consistent between the three isolates; however, there was a significant difference in the number of genes significantly up- or down-regulated under ciprofloxacin exposure, when compared to no treatment for D23580 (−2 ≥ log_2_fc ≥ 2, *p* <0.05; D23580: 259, SL1344: 165, VNS20081: 160) (**Figure 3, Supplementary data file S1**). Prophage and SOS response genes were amongst the most consistently highly upregulated regions in all isolates, and flagellar genes were most highly downregulated, although the number of genes and extent of upregulation was variable by isolate (**Table 2, 3**). Phage genes were not directly comparable between isolates, but in each isolate, they were the most highly upregulated genes, above SOS response genes (**Figure 3**). The top upregulated SOS response genes in common between the three isolates were *recN*, *sulA*, *recA*, *uvrA*, *lexA*, *sodA*, and *polB*, all genes known to be integral to the early bacterial stress response to double-stranded DNA damage (25–27). Interestingly, there were also several metabolism and biosynthesis-associated genes that were commonly upregulated (**Table 2**). Notably, other than flagellar genes, two downregulated genes in all isolates were *ompA* and *ompD*, which encode an outer membrane porin that plays a role in drug uptake and may be relevant in ciprofloxacin efflux (28–30). SL1344 had fewer downregulated genes with a log_2_ fold change ≤ −2, and the genes were less clustered along the chromosome than those of D23580 and VNS20081 (**Figure 3B**). **Table S1** and **Table S2** show the top 20 up- and down-regulated genes for *S.* Typhimurium D23580, respectively. The majority of upregulated genes were in prophage regions; downregulated genes were overwhelmingly associated with flagella and pili formation (**Figure 3A**). It is possible that D23580 had considerably more differentially expressed genes than SL1344 or VNS20081 because of a more robust prophage response.

**Table 2.** Top 20 significantly upregulated genes found commonly between SL1344, D23580, and VNS20081. *Differential expression analysis using DESeq2 was performed on each isolate independently for ciprofloxacin treatment versus no treatment, and only significant (adjusted *p-*value (padj) < 0.05) log_2_ fold change (l2fc) results were included. The top 20 upregulated genes for D23580 were sorted in descending order by l2fc and matched with corresponding l2fc for SL1344 and VNS20081. A padj value of “0” indicates the value was so small that it was rounded to 0 by DESeq2.

|  |  |  |  |  |  |  |  |
| --- | --- | --- | --- | --- | --- | --- | --- |
| *recN* | SOS response | 3.99 | 0 | 3.75 | 0 | 3.38 | 3.97E-134 |
| *sulA* | SOS response | 3.64 | 7.09E-108 | 3.54 | 2.1E-134 | 2.43 | 6.92E-22 |
| *recA* | SOS response | 3.31 | 0 | 3.46 | 0 | 2.66 | 1.78E-222 |
| *stdA* | Fimbriae production | 3.09 | 1.48E-06 | 2.68 | 3.74E-09 | 5.60 | 1.17E-14 |
| *ilvC* | Redox, biosynthesis | 2.01 | 4.71E-52 | 2.51 | 2.29E-115 | 2.15 | 1.17E-37 |
| *uvrA* | SOS response | 2.30 | 0 | 2.38 | 1.87E-288 | 1.93 | 1.21E-71 |
| *lexA* | SOS response | 2.26 | 9.71E-150 | 2.28 | 9.69E-257 | 1.89 | 3.11E-160 |
| *cysJ* | Redox, biosynthesis | 2.10 | 2.27E-35 | 2.26 | 3.3E-22 | 1.74 | 1.51E-15 |
| *cysD* | Redox, biosynthesis | 1.60 | 6.24E-09 | 2.17 | 6.27E-26 | 1.44 | 7.44E-09 |
| *cysH* | Redox, biosynthesis | 1.76 | 5.51E-13 | 2.13 | 6.95E-25 | 1.20 | 0.00002 |
| *leuA* | Biosynthesis | 2.02 | 2.15E-20 | 2.08 | 2.7E-61 | 1.87 | 2.01E-13 |
| *cysI* | Redox, biosynthesis | 1.77 | 3.62E-28 | 2.00 | 4.66E-26 | 1.42 | 5.28E-10 |
| *cysC* | Redox, biosynthesis | 1.46 | 5.35E-09 | 1.94 | 1.57E-23 | 0.89 | 0.019 |
| *sodA* | SOS response | 1.47 | 2.83E-71 | 1.94 | 1.19E-194 | 1.48 | 4.4E-27 |
| *fadB* | Redox, biosynthesis | 1.28 | 2.41E-06 | 1.92 | 1.58E-18 | 1.99 | 4.79E-10 |
| *cpxP* | Copper/H<sub>2</sub>O<sub>2</sub> resistance | 1.73 | 1.36E-24 | 1.86 | 2.44E-28 | 1.39 | 1.63E-14 |
| *polB* | SOS response | 2.27 | 2.24E-82 | 1.85 | 1.07E-42 | 1.80 | 1.13E-13 |
| *cysN* | Redox, biosynthesis | 1.65 | 9.53E-26 | 1.82 | 7.73E-29 | 0.85 | 0.0007 |
| *glmU* | Biosynthesis | 1.15 | 3.61E-110 | 1.80 | 3.87E-103 | 0.79 | 2.28E-23 |
| *fadA* | Redox, biosynthesis | 1.47 | 5.55E-06 | 1.79 | 9.25E-19 | 1.44 | 0.0007 |

**Table 3.** Top 20 significantly downregulated genes found commonly between SL1344, D23580, and VNS20081. *Differential expression analysis using DESeq2 was performed on each isolate independently for ciprofloxacin treatment versus no treatment, and only significant (adjusted *p-*value (padj) < 0.05)

| Gene* | Function | SL1344<br>l2fc | SL1344<br>padj | D23580<br>l2fc | D23580<br>padj | VNS20081<br>l2fc | VNS20081<br>padj |
| --- | --- | --- | --- | --- | --- | --- | --- |
| *flgH* | Flagellum | -2.17 | 2.16E-27 | -2.60 | 7E-80 | -2.52 | 7.39E-10 |
| *flgE* | Flagellum | -1.83 | 4.78E-30 | -2.58 | 1.06E-104 | -2.42 | 4.77E-12 |
| *flgJ* | Flagellum | -2.00 | 6.12E-21 | -2.53 | 2.42E-79 | -2.23 | 1.8E-15 |
| *flgF* | Flagellum | -2.00 | 2.90E-25 | -2.52 | 1.17E-77 | -2.47 | 9.24E-10 |
| *flgG* | Flagellum | -2.12 | 3.23E-33 | -2.49 | 5.77E-95 | -2.28 | 1.2E-09 |
| *flgI* | Flagellum | -1.97 | 1.54E-20 | -2.38 | 1.4E-62 | -2.31 | 6.41E-15 |
| *flgC* | Flagellum | -1.42 | 1.37E-17 | -2.37 | 3.11E-76 | -1.90 | 3.51E-08 |
| *flgB* | Flagellum | -1.67 | 7.61E-25 | -2.35 | 1.24E-61 | -2.06 | 1.11E-14 |
| *fliO* | Flagellum | -1.29 | 1.72E-11 | -2.33 | 3.05E-44 | -1.62 | 0.0000014 |
| *flgL* | Flagellum | -1.53 | 2.72E-20 | -2.28 | 4.34E-55 | -2.30 | 1.9E-24 |
| *yciH* | Putative translation factor | -1.38 | 8.11E-05 | -2.28 | 1.06E-22 | -1.66 | 0.0000188 |
| *flgK* | Flagellum | -1.44 | 4.35E-22 | -2.24 | 5.2E-98 | -2.40 | 1.42E-27 |
| *flgM* | Flagellum | -1.62 | 1.25E-14 | -2.07 | 1.58E-25 | -2.52 | 5.73E-18 |
| *yeeF* | Putative transporter | -1.46 | 4.07E-172 | -1.95 | 2.43E-242 | -1.43 | 4.36E-103 |
| *fliN* | Flagellum | -1.20 | 4.90E-10 | -1.92 | 2.43E-39 | -1.79 | 5.03E-12 |
| *fliI* | Flagellum | -0.98 | 1.41E-11 | -1.92 | 5.11E-09 | -1.87 | 1.86E-14 |
| *ybiN* | Conserved hypothetical protein | -1.80 | 5.83E-15 | -1.91 | 3.86E-39 | -1.16 | 0.000000294 |
| *fliL* | Flagellum | -1.11 | 1.67E-14 | -1.87 | 5.92E-46 | -1.47 | 0.00000177 |
| *flgN* | Flagellum | -1.59 | 2.12E-13 | -1.83 | 9.32E-21 | -1.88 | 2.25E-16 |
| *nth* | Biosynthesis | -1.52 | 6.14E-14 | -1.82 | 1.17E-32 | -1.96 | 5.07E-08 |

log<sub>2</sub> fold change (l2fc) results were included. The top 20 downregulated genes for D23580 were sorted in ascending order by l2fc and matched with corresponding l2fc for SL1344 and VNS20081. A padj value of “0” indicates the value was so small that it was rounded to 0 by DESeq2.

**Figure 3.**
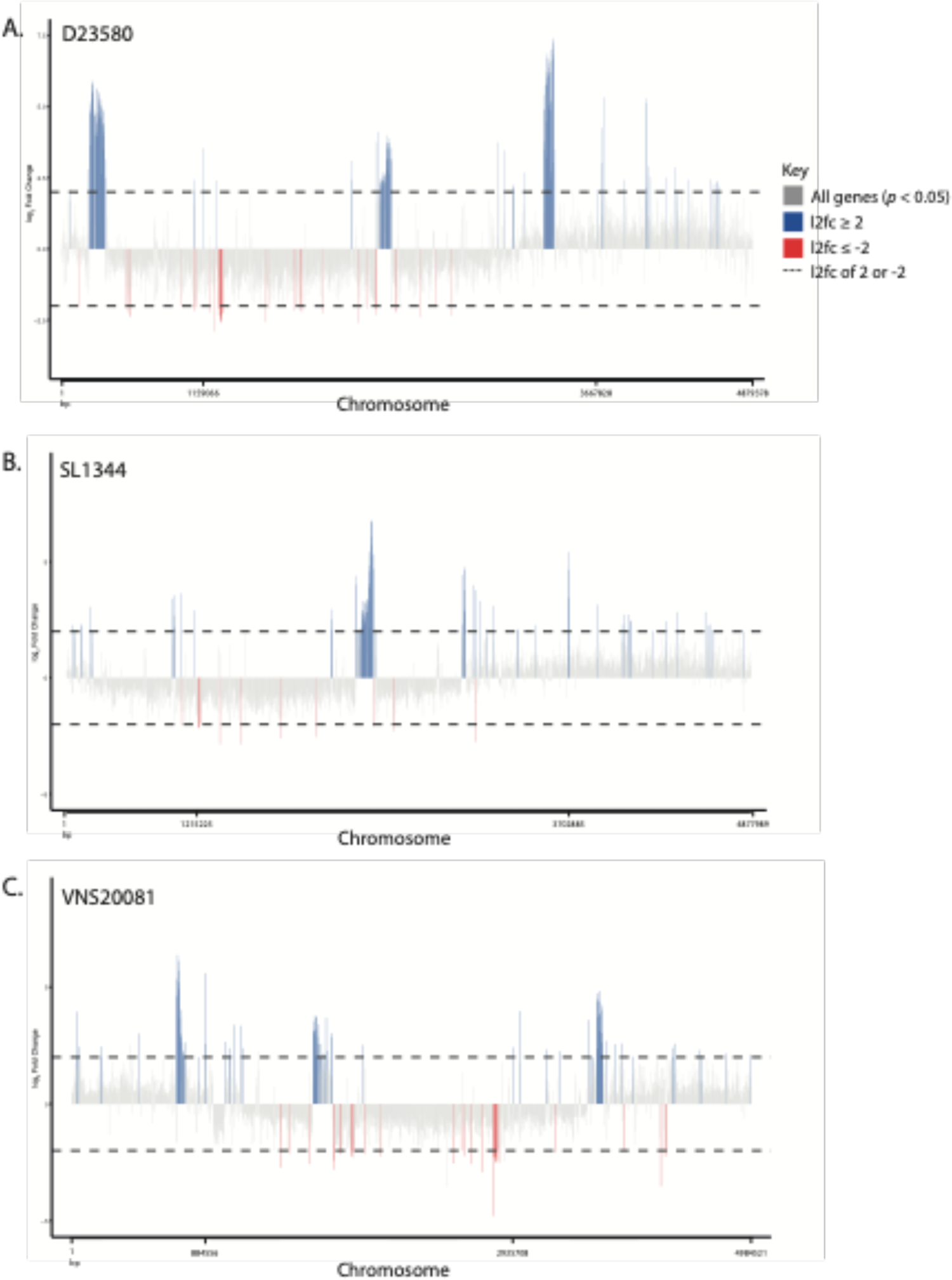
Bulk transcriptomics of *S.* Typhimurium following 2 hours of ciprofloxacin treatment. *S.* Typhimurium isolates D23580 (**A**), SL1344 (**B**), and VNS20081 (**C**) were grown in medium containing either 0× or 2× MIC ciprofloxacin for 2 hours, and RNA-sequencing was performed. Differential gene expression was analysed using DESeq2. The relative expression (log_2_ fold change) of each gene for 2× MIC ciprofloxacin versus 0× MIC ciprofloxacin was calculated for each isolate, and genes with an adjusted *p*-value < 0.05 were plotted along the chromosome. Genes with a log_2_ fold change ≥ 2 were coloured blue, and genes with a log_2_ fold change ≤ −2 were coloured red to highlight highly differentially-expressed genes.

### Ciprofloxacin triggers a specific dose dependant transcriptional response in D23580

To better understand the specificity of the bacterial stress response against ciprofloxacin, D23580 was subjected to four different perturbations: 0.5x and 2× MIC ciprofloxacin, Mitomycin C, and 1× MIC azithromycin, for two hours prior to RNA-sequencing. We chose D23580 to investigate further as it is an important clinical isolate for understanding invasive *Salmonella* disease and showed the most differential expression upon ciprofloxacin exposure. We found that each stressor induced a different transcriptional signature (**Figure 4, Supplementary data file S2**). We observed some overlap in transcriptional response between the two concentrations of ciprofloxacin, suggesting that a sub-inhibitory ciprofloxacin concentration (0.5x MIC) elicits a reduced SOS response (with respect to 2× MIC), although upregulation of prophage genes was comparable. Furthermore, there were fewer downregulated genes in the sub-inhibitory concentration of ciprofloxacin compared to 2× MIC (**Figure 4A–B**). In contrast, treatment with Mitomycin C, a potent inducer of double-stranded DNA breaks, elicited a notable prophage response (**Figure 4C**) (31–34), but fewer SOS response genes were upregulated than in the ciprofloxacin-treated conditions, and overall fewer genes were differentially expressed. This was surprising as we expected that Mitomycin C would elicit a similar transcriptional signature to ciprofloxacin; however, these differences may have been due to the concentration of Mitomycin C used. Such differences imply that while there is some overlap between the effects of ciprofloxacin and Mitomycin C, they are not identical, and ciprofloxacin elicits a distinct stress response. Most notably, treatment with azithromycin, an azide antimicrobial that targets the 50S ribosomal subunit, elicited a unique transcriptional profile, with no overlapping genes with any of the other conditions, indicating that the mode of action of the antimicrobial induces a specific impact on the transcriptional profile (**Figure 4D; Table S3, Table S4**). Importantly, this signified that the transcriptional response to inhibitory concentrations of ciprofloxacin is distinct from that to azithromycin, and this difference may be useful for considering treatment options in clinical settings.

**Figure 4.**
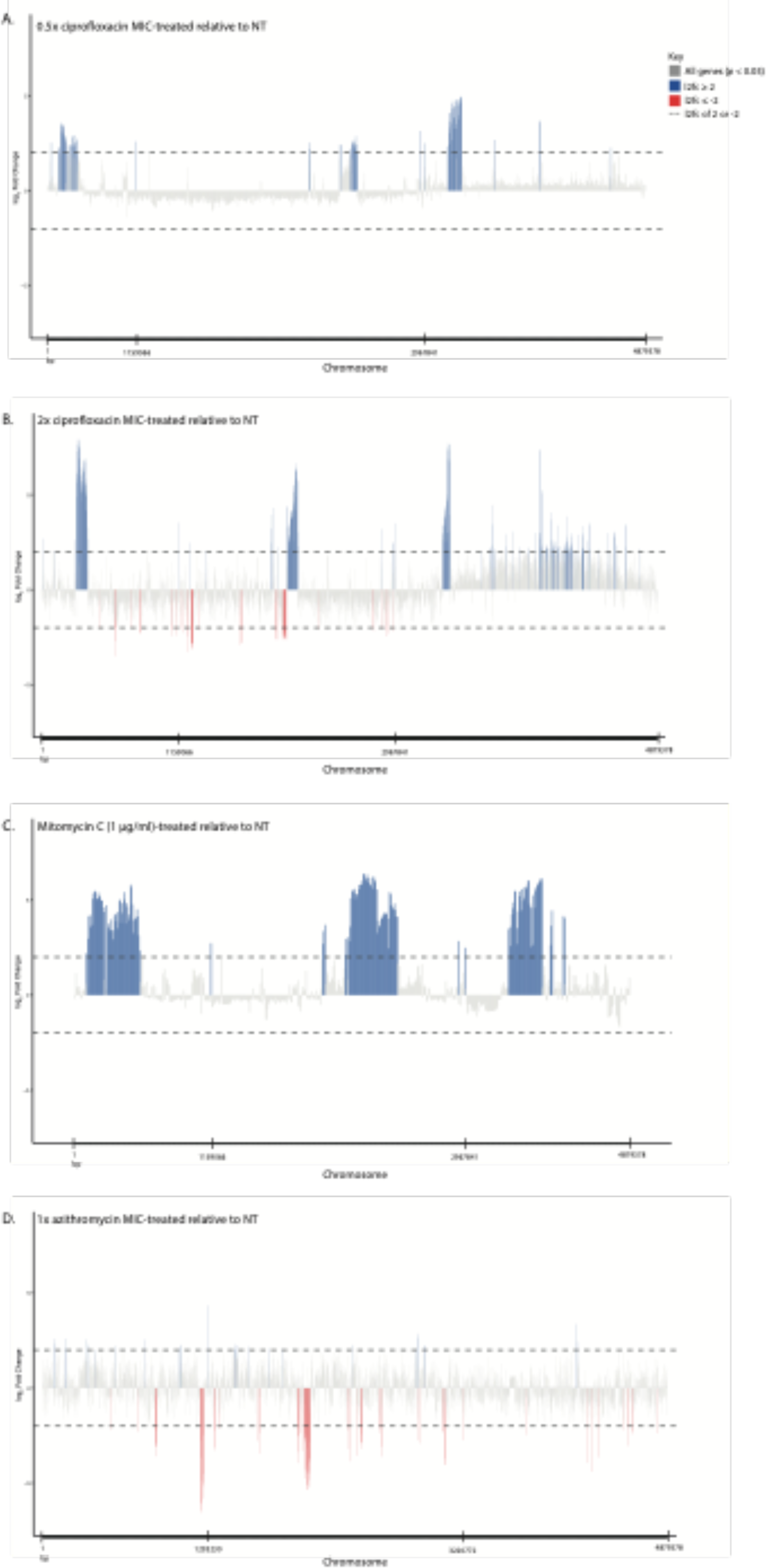
Bulk transcriptomics of *S.* Typhimurium D23580 under 4 different perturbations. *S.* Typhimurium D23580 was grown for 2 hours in medium containing 0.5x ciprofloxacin MIC (A), 2× ciprofloxacin MIC (B), 1 μg/ml Mitomycin C (C), or 1× azithromycin MIC (D) and subjected to RNA-sequencing. Differential gene expression was analysed using DESeq2. The relative expression (log_2_ fold change) of each gene for treatment versus no treatment was calculated for each condition, and genes with an adjusted p-value < 0.05 were plotted along the chromosome. Genes with a log_2_ fold change ≥ 2 were coloured blue, and genes with a log_2_ fold change ≤ −2 were coloured red to highlight highly differentially-expressed genes.

### Ciprofloxacin exposure stimulates a heterogeneous population with distinct transcriptional profiles

As demonstrated above, ciprofloxacin exposure induced pronounced morphological changes across the bacterial population. We wanted to determine whether these morphologically distinct bacteria could be physically separated and classified as subpopulations based on different physical and transcriptional properties. To disaggregate the transcriptional profiles associated with the various populations formed during ciprofloxacin exposure, we performed chilled sucrose density centrifugation of D23580 to separate elongated from non-elongated bacteria. The untreated D23580 bacteria formed a single diffuse fraction at approximately 50% sucrose, whereas the bacteria treated with 2× MIC ciprofloxacin segregated into three smaller fractions (within 50%, 60% and at the 60-70 % sucrose interface) (**Figure 5, Supplementary data file S3**). Based on our ability to separate morphologically distinct bacteria into specific fractions by density, we determined that there were meaningful subpopulations that formed in response to ciprofloxacin exposure. RNA sequencing of the three fractions generated with 2× MIC ciprofloxacin yielded markedly different transcriptional profiles. The low density (50% sucrose) and high density (60% sucrose) bacteria after 2× MIC ciprofloxacin exposure clustered independently with respect to their transcriptional profiles, which were also distinct from untreated bacteria (**Figure 5A; Tables S5-S8**).

**Figure 5.**
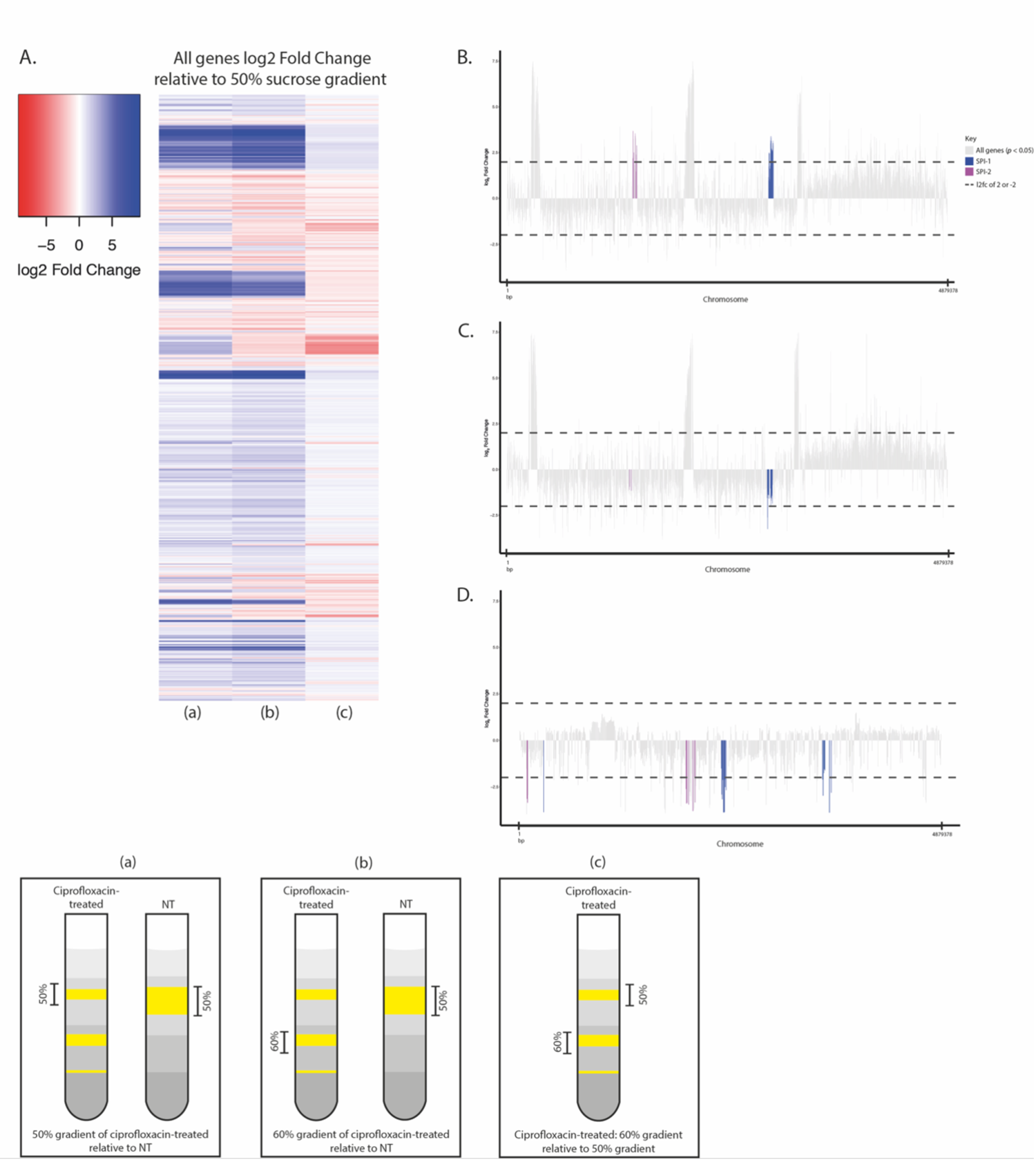
Transcriptomics of density gradient-separated *S.* Typhimurium D23580. *S.* Typhimurium D23580 was grown for 2 hours in either 0× (NT) or 2× MIC ciprofloxacin and layered on sucrose gradients containing 25%, 50%, 60%, and 70% sucrose layers. Following density centrifugation, gradient-separated bacteria were subjected to RNA-sequencing, and differential gene expression was analysed using DESeq2. **A.** Three comparisons were performed, and the log_2_ fold change of relative gene expression was plotted as a heatmap with upregulated genes in blue and downregulated genes in red. The comparisons were: ciprofloxacin-treated 50% sucrose gradient versus NT (a), ciprofloxacin-treated 60% sucrose gradient versus NT (b), and ciprofloxacin-treated 60% sucrose gradient versus ciprofloxacin-treated 50% sucrose gradient (c). **B.** For the comparison ciprofloxacin-treated 50% sucrose gradient versus NT, significantly differentially-expressed (p < 0.05) genes were plotted along the chromosome, and genes found within SPI-1 and SPI-2 were coloured in purple and blue, respectively. **C.** The comparison of ciprofloxacin-treated 60% sucrose gradient versus NT was mapped along the chromosome, as in B. **D.** The comparison of ciprofloxacin-treated 60% sucrose gradient versus ciprofloxacin-treated 50% sucrose gradient, as in B.

An analysis of the top upregulated and downregulated genes showed that >100 genes were downregulated in the high density compared to the low-density ciprofloxacin-treated bacteria (**Figure 5B, (c)**). Specifically, fewer genes were upregulated in the high-density fraction in comparison to the low-density fractions of the ciprofloxacin-treated bacteria. We observed that Salmonella Pathogenicity Island 1 (SPI-1) and other invasion-associated genes were downregulated in the high-density ciprofloxacin-treated bacteria. In contrast, there was significant upregulation of some SPI-1 and SPI-2 genes in the low density (50% sucrose) ciprofloxacin-treated bacteria compared to untreated bacteria (**Figure 5 B–C; Tables S5-S8**). These data suggest that under ciprofloxacin treatment, elongated bacteria suppress genes that trigger cellular invasion and have an elevated stress response compared to non-elongated bacteria; additionally, the data suggests that ciprofloxacin-treated non-elongated bacteria may be better primed for cellular invasion and replication.

### Ciprofloxacin exposure impacts host-pathogen interactions

We observed that ciprofloxacin-exposed D23580 downregulates SPI-1 and SPI-2 genes, suggesting that ciprofloxacin may impact on the ability of *S.* Typhimurium to invade and replicate in host cells. Therefore, we tested this hypothesis by assessing the interaction between ciprofloxacin exposed D23580 with macrophages and epithelial cells using a modified gentamicin protection assay (35, 36). To this end, bacteria were cultured for two hours in the absence or presence of 0.06 μg/ml ciprofloxacin and subsequently inoculated onto monolayers of macrophages or HeLa cells. At 1.5 hours post-infection, a significantly larger percentage of the ciprofloxacin-treated inoculum was internalized by macrophages (mean % internalized of inoculum: 6.72% versus 1.50%, *p* < 0.005) (**Figure 6A left panel**). This difference is significant given that the inoculum added to cells, as measured by CFU/ml, was 100-fold lower for the ciprofloxacin-treated bacteria given two hours of ciprofloxacin exposure (2.83E+06 ± 1.15E+06 CFU/ml for untreated bacteria versus 1.63E+04 ± 4.62E+03 CFU/ml for ciprofloxacin-treated bacteria). Thus, although significantly fewer ciprofloxacin-treated bacteria were added to equivalent numbers of macrophages, a significantly higher percentage of treated bacteria were internalized. Furthermore, the ciprofloxacin-treated bacteria had a higher replication rate in macrophages than untreated bacteria at 6 hours post-infection (mean fold replication over 1.5 hours: 0.66 versus 0.20, *p* < 0.05) (**Figure 6A right panel**). It is possible that the macrophages internalized ciprofloxacin-treated bacteria at a higher rate because of their increased size and lower viability. However, this did not explain the greater intracellular survival and fold replication of ciprofloxacin-treated bacteria within macrophages.

**Figure 6.**
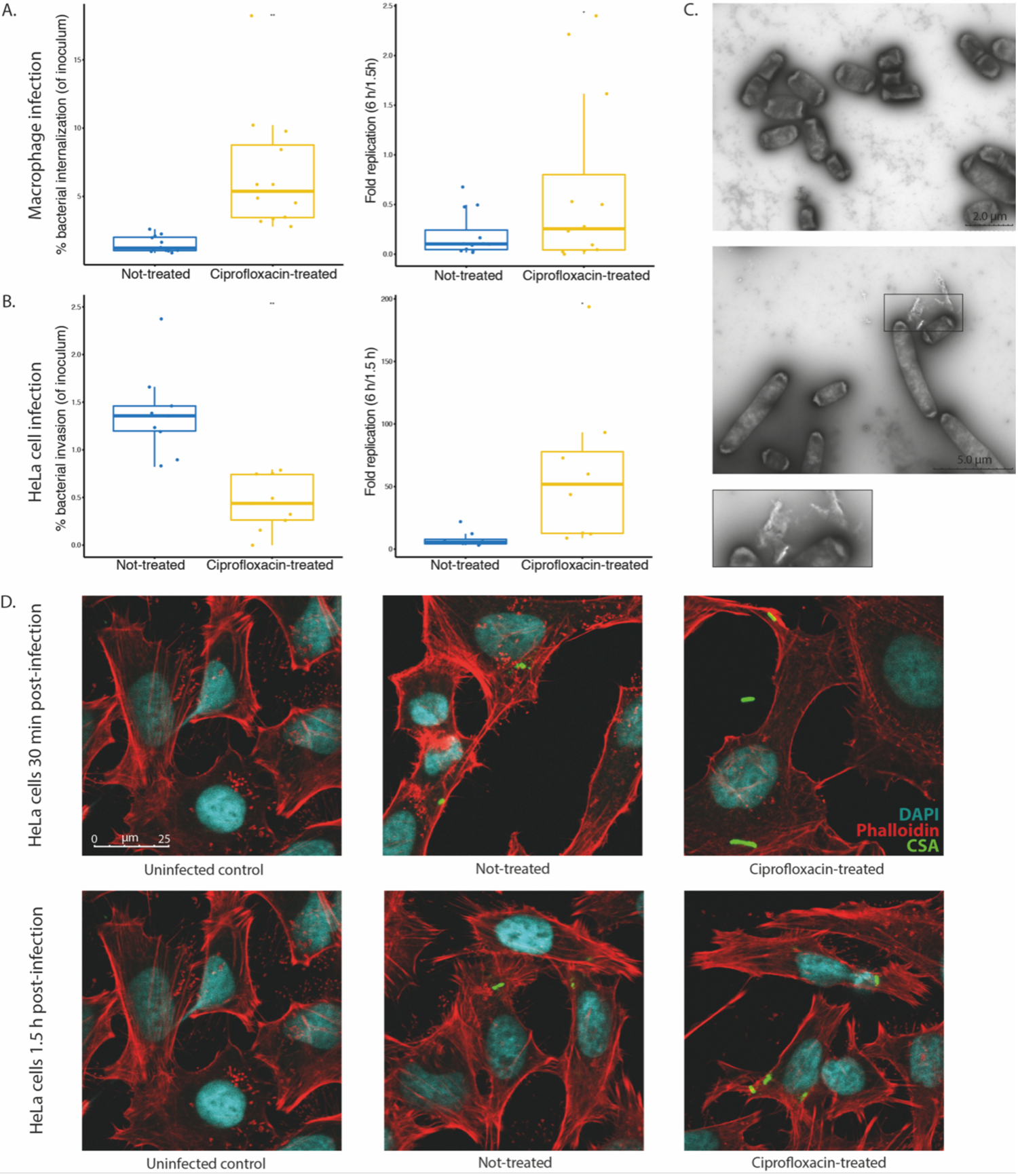
Cellular infections with *S.* Typhimurium D23580 following 2 hours of ciprofloxacin exposure. *S.* Typhimurium D23580 was either not treated or treated with 2× MIC ciprofloxacin for 2 hours prior to infection of macrophages (**A**) or HeLa cells (**B**). Left panels show bacterial internalization 1.5 hours post-infection. Right panels show bacterial intracellular replication 6 hours post-infection. Boxplots represent the mean and interquartile ranges of four (macrophages) or three (HeLa cells) biological replicates of three technical replicates each. The mean and SD were calculated, and a Student’s paired t-test was performed to calculate significance. * *p* < 0.05. **C.** Transmission electron microscopy was performed using negatively-stained D23580 either not treated (top panel) or treated with 2× MIC ciprofloxacin (bottom panel) for 2 hours. Box inset shows extracellular matter in ciprofloxacin-treated culture. **D.** Confocal images were taken of D23580 either not treated or treated with 2× ciprofloxacin MIC immediately following the initial 30 minutes infection of HeLa cells (top panel) or after the subsequent one-hour gentamicin treatment (bottom panel). HeLa cell membranes were stained with phalloidin (red), nucleic acids were stained with DAPI (blue), and bacteria were stained with CSA (green). Images of infected cells are compared to an uninfected control image for reference (left panel, same image used as comparator for 30 minutes and 1.5 h).

To investigate whether the ciprofloxacin-treated bacteria were actively modulating interactions with host cells, the same assay was repeated using HeLa cells. We found that ciprofloxacin-treated D23580 bacteria displayed significantly lower rates of infection than untreated bacteria (mean % internalized of inoculum: 0.44 % versus 1.38%, *p* < 0.005) (**Figure 6B left panel**). However, in a comparable manner to macrophages, the fold replication 6 hours post-infection of ciprofloxacin-treated bacteria was significantly higher than that of the untreated bacteria (mean fold replication over 1.5 hours: 62.14 versus 7.88, *p* < 0.05) (**Figure 6B right panel**). This observation suggests that ciprofloxacin exposure diminishes invasion rates of epithelial cells, but makes intracellular replication more efficient.

We hypothesized that debris from bacteria killed by ciprofloxacin may influence bacterial uptake by host cells. To assess whether the cultures of untreated and ciprofloxacin-treated bacteria differed, transmission EM was performed on the two cultures prior to infection (**Figure 6C**). Using a negative stain, we observed that some ciprofloxacin-treated bacteria appeared to be associated with extracellular matter of unknow origin (**Figure 6C lower panel, inset**). We could not identify substantial differences in the cultures of untreated and ciprofloxacin-treated bacteria, but further study may be warranted to determine whether ciprofloxacin-killed bacteria influence survival of live bacteria in the same environment.

To determine whether the bacterial morphology influenced invasion of HeLa cells, bacteria were imaged immediately following the 30-minute infection and at 1.5 hours, following 1 hour gentamicin treatment (**Figure 6D**). At 30 minutes post-infection, there were elongated and non-elongated ciprofloxacin-treated bacteria extracellularly (**Figure 6D top right panel**). In contrast, imaging at 1.5 hours post-infection showed similar sized and shaped untreated and ciprofloxacin-treated internalised bacteria in HeLa cells (**Figure 6D bottom row**). Our data suggest that non-elongated ciprofloxacin-treated bacteria are more efficient than elongated ciprofloxacin-treated bacteria at invading HeLa cells, and it is possible that invasion by this subpopulation may enhance intracellular survival and replication.

## Discussion

Here, we investigated morphological and transcriptional responses of three distinct *S.* Typhimurium isolates against measured inhibitory concentrations of ciprofloxacin. We found that these bacteria were highly resilient to increasing concentrations of ciprofloxacin and adapt to this environment over a 24-hour period of antimicrobial exposure, forming morphologically and transcriptionally distinguishable subpopulations early after exposure that have enhanced capacity to invade cells and replicate. Importantly, these data better define how clinical isolates respond to ciprofloxacin exposure, illustrating the potential for clinical *S.* Typhimurium isolates to tolerate and even replicate in the presence of concentrations of ciprofloxacin that should be fatal.

Ciprofloxacin (and other fluoroquinolones) are known to upregulate the bacterial stress response and phage activity (confirmed here) and the widespread use of ciprofloxacin is likely exacerbating AMR (18, 37). While population heterogeneity has been observed in response to ciprofloxacin exposure, past studies have used sub-inhibitory concentrations of ciprofloxacin against *E. coli* (38, 39). Our work shows that sub-inhibitory concentrations of ciprofloxacin have a muted effect on the bacterial response and may be less relevant for understanding the bacterial response to clinical dosages. However, previous observations are in concordance with our findings that bacterial subpopulations have highly distinct transcriptional responses, which may imply a bet-hedging strategy to improve survival potential. Importantly, we also showed that the response to ciprofloxacin is specific and dosage-dependent, and the upregulation of stress response and error-prone DNA replication machinery may influence bacterial survival and mutation (40–44). One limiting aspect of our study was that we did not longitudinally follow the bacterial response to ciprofloxacin, and future studies should also explore whether the ciprofloxacin MIC changes within a short time frame, and how that affects the transcriptional response.

While not explored in this study, other groups have studied bacterial persistence in relation to ciprofloxacin at length (45–49). Bacterial persistence may factor into observations made in this study; however, one critical difference is that bacteria were consistently, rather than intermittently, exposed to ciprofloxacin. The ability of the bacteria to grow under constant ciprofloxacin pressure and subsequently invade host cells suggests additional factors are involved in cellular survival and resilience to ciprofloxacin during exposure.

Our work additionally suggests that ciprofloxacin-treated bacteria have somewhat different infection dynamics compared to untreated bacteria, which may have broader implications for patients on fluoroquinolone treatment. The invasion of, and replication within, HeLa cells and macrophages of *S.* Typhimurium has been well-characterized, and many pathways involved in efflux and drug resistance have also been studied in the context of host-pathogen interactions (50, 51). Work by Anuforom *et al.* found that J774 murine macrophages expressed greater concentrations of IL-1ß and TNF-α when pre-treated with ciprofloxacin in the presence of SL1344. Additionally, they observed greater bacterial adhesion to ciprofloxacin-treated macrophages, resulting in enhanced bacterial killing (52). One limitation of our study is that we did not compare bacteria pre-treated with ciprofloxacin to those exposed to ciprofloxacin within host cells. Given the findings of Anuforom *et al.* and the importance of intracellular survival, intracellular interactions with ciprofloxacin may play a key role in drug evasion, and future work should investigate the response of *S.* Typhimurium to ciprofloxacin after cellular internalization.

However, in our study, we focused on the extracellular impacts of ciprofloxacin exposure, and the influence of ciprofloxacin treatment on bacteria prior to the infection of epithelial cells and macrophages has not been extensively studied. While we observed differences in the infection and replication potential between ciprofloxacin-treated and untreated *S.* Typhimurium that associated with transcriptional changes occurring in bacterial subpopulations, we did not investigate specific loci that could be responsible for the observed phenotype. It would be valuable to investigate any potential role in ciprofloxacin escape at the gene level to better understand how ciprofloxacin treatment may further affect *Salmonella* infections.

In a climate of mass drug administration (MDA) in parts of the world, it is particularly important to be aware of and actively study how bacteria respond to widespread antimicrobial exposure. In recent years, MDA studies have included single-dose administration of ciprofloxacin to combat *Neisseria meningitidis* in young children in the “meningitis belt” of Africa, prophylactic azithromycin in Niger, Malawi, and Tanzania to reduce childhood mortality, and azithromycin administration for children with non-bloody diarrhoea in low resource settings (53–55). While initial follow-up studies into resulting AMR have been performed, more genotypic and phenotypic surveillance is required (56). The potential for ciprofloxacin to trigger adaptive and genetic resistance in bacteria that may improve bacterial survival intracellularly provides impetus for greater caution in fluoroquinolone usage and more detailed investigation of the effect of ciprofloxacin and other antimicrobials on host-pathogen interactions.

## Materials & Methods

### Bacterial isolates and growth medium

Three *Salmonella* Typhimurium isolates were used: SL1344 (ST19, United Kingdom), VNS20081 (ST34, Vietnam), and D23580 (ST313, Malawi) (22, 23, 57). Prior to experimentation, all isolates were grown on Isosensitest agar (Oxoid, CM0471) and subjected to M.I.C.E. (Oxoid, MA0104F) ciprofloxacin eTests in duplicate to determine baseline ciprofloxacin susceptibility and MIC range was confirmed by assessment on the Vitek2 (**Table 4**). Isolates were grown in Isosensitest broth (Oxoid, CM0473) for all except host cell experiments and were maintained on Isosensitest agar and streaked weekly from frozen stocks.

**Table 4.** MICs using Vitek 2 and ciprofloxacin eTest.

|  | Vitek2 ciprofloxacin result | M.I.C.E. ciprofloxacin eTest result<br>(µg/ml) |
| --- | --- | --- |
| **D23580** | <=0.25 | 0.03 |
| **SL1344** | <=0.25 | 0.015 |
| **VNS20081** | 2 | 1 |

### Time kill curves

Colonies from plates were inoculated in 10 ml Isosensitest broth for 16-18 h shaking at 200 rpm at 37°C. Bacteria were added in a 1:10000 dilution to 10 ml of Isosensitest containing levels (0×, 1×, 2×, 4× MIC) of ciprofloxacin according to each isolate’s MIC for an inoculum of between 1 and 5 × 105 CFU/ml. Cultures were incubated shaking at 37°C and aliquots were taken for CFU plating at 0, 2, 4, 6, 8, and 24 h. Serial dilutions were made, and a total of 50 μl of each dilution was plated in 10 μl on L-agar. CFUs were counted and calculated as CFU/ml. Mean and SD of three replicates per isolate were calculated. Log_10_ CFU/ml were plotted over 24 hours as three independent replicates, with the colour indicating growth condition (0×, 1×, 2×, 4× ciprofloxacin MIC) in R using ggplot2 (58, 59). To compare mean CFU/ml for the 24-hour timepoint, an analysis of variance (ANOVA) was performed, and statistical significance of differences in the means of conditions compared to 0× (control) were conducted using Dunnett’s test.

### Ciprofloxacin-degradation kill curves

Initial 24 h time kill curves were performed as described above. At 24 h, cultures were centrifuged and steri-filtered, and filtered medium was transferred to fresh tubes. As above, overnight cultures were added 1:10000 to the medium and CFU were plated at 0, 2, 4, 6, 8, and 24 h. No additional ciprofloxacin was added to medium.

### RNA extractions and RNA sequencing

After bacteria were subcultured 1:1000 for 2 h in the presence or absence of 2× ciprofloxacin MIC, double the quantity of RNAProtect Bacteria Reagent (Qiagen, 76506) was added to cultures and incubated for 10 min. Cultures were centrifuged at 3215 × *g* for 14 min at 4°C. Supernatant was decanted and resuspended in 400 μl Tris buffer (0.25 mM, pH 8.0) containing 10 mg/ml lysozyme, and incubated for 5 min. To this was added 700 μl RLT buffer containing 10 μl beta-mercaptoethanol (Sigma, M6250) per ml and vortexed well. 1 ml 100% ethanol was immediately added and vortexed well. The Qiagen RNeasy Mini Kit (Qiagen, 74104) was subsequently used to process samples. Samples were eluted in 40 μl RNase-free water. Samples were frozen at −20°C if not immediately processed. Subsequently, samples were treated with DNase I using the Qiagen DNase Kit (Qiagen, 79254). Output following DNase treatment was cleaned using phenol-chloroform treatment by first increasing solution volume with RNase-free water to 400 μl. 400 μl of phenol-chloroform-isoamyl alcohol mixture (Sigma, 77617) was added to samples, mixed by inversion, then centrifuged at 8000 × *g* for 5 min. The supernatant was transferred to a new tube and combined with 400 μl chloroform:isoamyl alcohol 24:1 (Sigma, C0549). Samples were mixed then centrifuged as above. The supernatant was transferred to a new tube and combined with 1 μl glycogen (Roche, 10901393001), 40 μl 3M sodium acetate, pH 5.5 (Ambion, AM9740), and 500 μl ice-cold 100% ethanol. Tubes were mixed by inversion and incubated at −20°C for 30 min before centrifugation at 4°C for 20 min at 16000 × *g*. Supernatant was decanted and replaced with 500 μl ice-cold 70% ethanol and centrifuged at 4°C for 5 min at 16000 × *g*. Ethanol was decanted and pellets were air-dried before resuspension in 50 μl RNase-free water. Samples were frozen at −80°C prior to sequencing. All library preparation and RNA-sequencing were performed at the Wellcome Sanger Institute using standard protocols. Briefly, libraries were made using the NEB Ultra II RNA custom kit (NEB, E7530S) on an Agilent Bravo WS automation system. RiboZero was added to deplete ribosomal RNA. Libraries were pooled and normalized to 2.8 nM for sequencing. Sequencing was done on an Illumina HiSeq 4000 (Illumina, San Diego, CA), using a minimum of two lanes per pool.

### RNA sequencing analysis

Reads from D23580 were mapped to reference sequence D23580 (accession number <u>FN424405.1</u>) (22), VNS20081 was mapped to sequence VNB151 (accession number <u>ERS745838</u>) (23), and SL1344 was mapped to reference sequence SL1344 (accession number <u>FQ312003.1</u>). Sanger Institute pipeline DEAGO (Differential Expression Analysis & Gene Ontology), a wrapper script for DESeq2 and topGO (60, 61), was used to determine differential gene expression. Using DESeq2, a Wald test was done on the treatment condition versus untreated. The log_2_ fold change was calculated for treatment condition versus untreated after filtering genes to include only those with an adjusted *p*-value (padj) of < 0.05 to control for the false discovery rate using the Benjamini-Hochberg procedure. All differential expression analyses were conducted using default DESeq2 parameters (60). Genes that had a padj < 0.05 and a log_2_ fold change of ≥ 2 or ≤ −2 were subjected to further manual analysis to assess top up- and down-regulated genes in treatment conditions relative to untreated. Visualization of differentially-expressed genes was performed using the ggplot2 package in R.

### Sucrose gradient separation of ciprofloxacin-treated D23580

To separate morphologically distinct subpopulations of bacteria after ciprofloxacin exposure, a sucrose gradient procedure was developed. Overnight cultures of D23580 were grown as described above and were inoculated 1:100 into 10 ml of Isosensitest broth either containing 0 or 0.06 μg/ml ciprofloxacin and incubated shaking at 200 rpm at 37°C for 2 h. Fresh sucrose solutions were prepared: the four concentrations of sucrose used were 25%, 50%, 60%, and 70%, and these were made by dissolving sucrose (Sigma, S7903) in 1× PBS. Solutions were sterile-filtered using 0.2 μm syringe filters (GE Healthcare, 6794-2502). 2 ml of each sucrose concentration was layered from 70% to 25% in open-top ultracentrifuge tubes (Beckman Coulter, 344059) immediately before use. At 2 h, cultures were removed from incubator and centrifuged in a benchtop swing bucket centrifuge for 14 min at 4000 × *g* at 4°C. The supernatant was removed with a pipette. Pellets were resuspended in the remaining medium and transferred to 1.5 ml tubes, which were centrifuged at 5000 × *g* for 2 min to re-pellet. The supernatant was removed, and pellets were resuspended in 500 μl PBS. Using a Pasteur pipette, 500 μl of cells was carefully added to the top of the 25% layer of the sucrose column. Gradients were centrifuged for 9 min at 3000 × *g*, 4°C. After centrifugation, gradients were identified:

- One layer on the gradients loaded with non-treated cultures within the 50% sucrose fraction.
- Three layers from the 2× MIC ciprofloxacin-treated gradients: 1. within 50%, 2. within 60%, 3. 60-70% interface

The cloudy portion of each layer was carefully removed using a Pasteur pipette, beginning with the lowest-density layer, and isolated fractions were immediately added to 10 ml bacterial RNAProtect and processed using the standard RNA extraction protocol described above.

### RNA sequencing analysis of gradient-separated bacteria

RNA-seq analysis was performed on the bacteria recovered from the gradients. These RNA-sequencing reads were processed using DEAGO. Pairwise comparisons were made between conditions (ciprofloxacin-treated 50% vs untreated 50%, ciprofloxacin-treated 60% vs untreated 50%, and ciprofloxacin-treated 60% vs ciprofloxacin-treated 50%). Heatmaps were made using the heatmap.2 function in R package gplots, and other visualizations were performed using ggplot2.

### DNA extraction of 24 h ciprofloxacin-treated cultures

To prepare DNA, bacterial cultures of *S.* Typhimurium D23580 were initially grown overnight in 10 ml of broth. As in the time kill curve experiments, 10 ml of fresh Isosensitest broth containing no or 0.06 μg/ml ciprofloxacin MIC were inoculated with overnight cultures at 1:10000. Bacteria were grown for 24 h before spreading 100 μl or 1000 μl for the untreated and ciprofloxacin-treated cultures, respectively, on L-agar plates. Plates were grown overnight to ensure only DNA from viable organisms was sequenced as a plate sweep (62). After overnight growth at 37°C, colonies were scraped from the agar and resuspended in 1× PBS. This was spun down at 8000 rpm for 3 min, and the supernatant was aspirated off. The pellets were processed for DNA extraction using the Promega Wizard DNA Purification kit (Promega, A1120). DNA was quantified on a Qubit 4 Fluorometer (Q33226) using the Qubit dsDNA HS Assay Kit (Q32851), then frozen at −80°C prior to whole genome sequencing. DNA was sequenced on an Illumina HiSeq platform. Illumina adapter content was trimmed from reads using Trimmomatic v.0.33.

### Read mapping and variant detection of 24 h ciprofloxacin-treated cultures

Illumina HiSeq reads were mapped to *S.* Typhimurium reference genome D23580 (<u>FN424405.1</u>) using SMALT v0.7.4 to produce a BAM file. Briefly, variant detection was performed as previously detailed (63). Samtools mpileup v0.1.19 with parameters -d 1000-DSugBf and bcftools v0.1.19 were used to generate a BCF file of all variant sites. The bcftools variant quality score was set as greater than 50, mapping quality was set as greater than 30, the allele frequency was determined as either 0 for bases called same as the reference or 1 for bases called as a SNP (af1 < 0.95), the majority base call was set to be present in at least 75% of reads mapping at the base (ratio < 0.75), the minimum mapping depth was four read, a minimum of two of the four had to map to each strand, strand_bias was set as less than 0.001, map_bias less than 0.001, and tail_bias less than 0.001. Bases that did not meet those criteria were called as uncertain and removed. A pseudo-genome was constructed by substituting the base calls in the BCF file in the reference genome. Recombinant regions in the chromosome such as prophage regions were removed from the alignment and checked using Gubbins v1.4.10. SNP sites were extracted from the alignment using snp-sites and analysed manually.

### Opera Phenix confocal microscopy phenotyping of single bacteria

*S.* Typhimurium D23580, SL1344, and VNS20081 were screened at 2 h after ciprofloxacin exposures of 0×, 1×, 2×, and 4× as related to the MIC of the isolate. This was undertaken by inoculating overnight cultures independently at 1:1000 dilutions of 150 μl in 150 ml Isosensitest broth in a 200 ml flask incubated shaking. Following 2 h growth, 10 ml of each culture were spun down at 3200 × *g* for 7 min at 4°C. The supernatant was decanted, and the pellet was transferred to a 1.5 ml tube. This was spun at 8000 × *g* for 3 min, and the supernatant was decanted and replaced with 100 μl PBS. For each culture condition, 50 μl of the concentrated bacterial culture was added to two wells of a vitronectin-coated Opera CellCarrier Ultra-96 plate (Perkin Elmer, 6055302), and the plates were incubated static at 37 °C for 10 min. The microbial culture was aspirated, then fixed with 4% PFA, and washed with 1× PBS. Wells were incubated with 2% BSA for 30 min, then for 1 h with CSA-Alexa-647 (Novus Biologicals, NB110-16952AF647) at 1:1000 in BSA. Wells were aspirated and then incubated with solutions harbouring DAPI (Invitrogen, D1306) and SYTOX green (Invitrogen, S7020) for 20 min. Wells were washed 1× with PBS; plates were sealed and imaged.

### Opera Phenix confocal microscopy image analysis of single bacteria

Images generated on the Opera Phenix were analysed using the Harmony software (Perkin Elmer), as previously described (24) (S. Sridhar and S. Forrest, submitted for publication). Briefly, inputted images underwent flatfield correction, and images were calculated using the DAPI and Alexa647 channels and then refined by size and shape characteristics. Applying a linear classifier to the filtered population, single bacteria were identified, and morphology and intensity characteristics were calculated. The output of the Harmony analysis was tabulated by object, and results were visualized in R (v 3.6.1) using R packages dplyr and ggplot2.

### HeLa cell and iPS macrophage infections with S. Typhimurium D23580

HeLa cells were obtained from Abcam (ab255928) and maintained in DMEM + (Thermo, 41966) supplemented with 10% heat-inactivated FBS (Merck, F7524) incubated at 37°C, 5% CO2. HeLa cells were plated in 24-well plates (Corning, 3473) at 1 × 10^5^ cells/ml in 500 μl media. D23580 was inoculated from a freshly streaked plate in 10 ml LB and incubated shaking overnight at 37°C the day prior to infections. On the day of infections, two D23580 sub-cultures were set up 1:10 in LB from the overnight culture, with one sub-culture containing 0.06 μg/ml of ciprofloxacin. Cultures were incubated shaking at 37°C for 2 h. At 2 h, the OD600 of cultures was measured, and bacteria were resuspended in PBS after normalization to an OD600 of 1.0. Bacteria were added to cell media for a multiplicity of infection of ~10:1. 500 μl of the inoculum was added to each well and incubated for 30 min. The inoculum was plated for CFU enumeration. Following the infection, media was aspirated and cells were washed 1× with PBS. PBS was replaced with media containing 16 μg/ml gentamicin (Gibco, 15750037), and plates were incubated for 1 h. Media was aspirated and plates were washed 1× with PBS and subsequently either replaced with 0.1% Triton-X for the 1.5 h time point or media until 6 h post-infection. To enumerate CFU, 100 μl of cell lysates was spread on L-agar plates, and plates were incubated overnight at 37°C before counting. The same process was followed at 6 h post-infection. Infections were conducted in technical triplicates.

Macrophages derived from induced pluripotent stem cells were produced as previously described (Alasoo et al., 2015). Monocytes in RPMI containing hMCSF cytokine (Bio Techne/216-MC-025) were plated in 24-well plates at 1.5 × 10^5^ 7 days prior to infection, and the media was changed to RPMI without hM-CSF one day prior to infection. Cells were infected with D23580 as described above for HeLa cells, and CFU were enumerated.

### Confocal microscopy of infected HeLa cells

1 × 10^5^ HeLa cells/ml were added to coverslips (Thermo, 12392128) in 24-well plates, and infections with D23580 were conducted as above. After the 30 min infection, one set of coverslips were immediately fixed in 4% PFA without washing to image intracellular and extracellular bacteria. The remaining coverslips were processed as CFU wells and fixed at 1.5 h post-infection. Coverslips were blocked and permeabilized using 250 μl 10% BSA + 0.1% Triton X-100 in PBS for 15 minutes at room temperature. CSA (BacTrace, 5330-0059) and phalloidin (A22287) antibodies were diluted in 1% BSA + 0.1% Triton X-100 in PBS at 1:100 and 1:1000, respectively. 250 μl of the CSA antibody was added first and incubated in the dark at room temperature for 1 h. Coverslips were washed 3× in 250 μl PBS, and then 250 μl of phalloidin was added to coverslips and incubated in the dark at room temperature for 1 h. Coverslips were washed 3× in 250 μl PBS. Coverslips were mounted on glass slides with 20 μl Prolong Gold with DAPI (Invitrogen, P36935) and cured in the dark at room temperature overnight. 25 fields per coverslip were imaged on a Leica TCS SP8 confocal microscope at 40× magnification.

### Transmission electron microscopy (TEM) of S. Typhimurium D23580

D23580 overnight cultures were added 1:10 to 10 ml LB either containing none or 0.06 μg/ml ciprofloxacin and incubated shaking for 2 h. For staining, 1 ml of uranyl acetate (UA) solution (3% aqueous) was filter-sterilized through a 0.2 μm filter. One 200 square mesh Cu EM grid (Agar Scientific) was spotted with 10 μl bacterial sample and left for 1 min. Filter paper was used to remove excess liquid, and 10 μl UA was added to the grid for 1 min. Excess liquid was again removed using filter paper, and the grid was allowed to dry for 1 h prior to imaging. Imaging was done on a Hitachi HT7800 transmission electron microscope at 100kV, 8μA, and a range of magnifications.

## Data availability

RNA-sequencing reads can be found using the study accession number <u>PRJEB43116</u> (ERP127047). Whole genome sequencing reads can be found using the accession number <u>PRJEB43255</u> (ERP127204). **Supplementary data file S4** matches read files with samples. All supplementary data files can be found at https://doi.org/10.17605/OSF.IO/N9CW5.

## Acknowledgements

This work was supported by Wellcome (grant 206194) and the Wellcome Sanger Institute (PhD studentship to SS). SB is supported by a Wellcome senior research fellowship (215515/Z/19/Z). This work was supported by a Innovate UK Commercial in Confidence grant to purchase the Opera Phenix. SF, SB, CC, DP, and GD are supported by funding from the National Institute for Health Research [Cambridge Biomedical Research Centre at the Cambridge University Hospitals NHS Foundation Trust] and National Institute for Health Research AMR Research Capital Funding Scheme [NIHR200640]. *The funders* had *no role* in the design and conduct of the study; collection, management, analysis, and interpretation of the data; preparation, review, or approval of the manuscript; and decision to submit the manuscript for publication. We are grateful to Sina Beier for help with the RNA-sequencing analysis and Sandra Van Puyvelde for helpful discussions during the project. We are further grateful to the Wellcome Sanger Institute Sequencing Pipelines team for sequencing assistance and to the Wellcome Sanger Institute Pathogen Informatics team for bioinformatics support.

## Supplementary figures

**Figure S1. Time kill curves of *S.* Typhimurium under ciprofloxacin exposure to assess ciprofloxacin stability A.** Time kill curves were performed on *S.* Typhimurium D23580 using spent medium following an initial 24-hour kill curve. Media for this growth curve was centrifuged and steri-filtered before inoculation with D23580 and growth over 24 hours. CFU were enumerated at 6 time points, and two independent biological replicates were plotted. **B.** Average CFU/ml were plotted as mean ± SD for the 24-hour time point to compare CFU between treatment conditions. An ANOVA was performed to compare means at 24 hours, and Dunnett’s test was performed to compare 24 hour means of 1×, 2×, and 4× ciprofloxacin MIC to 0× (control).

## Supplementary tables

**Table S1. Top 20 significantly upregulated genes in 2× MIC ciprofloxacin-treated D23580 relative to NT.**

**Table S2. Top 20 significantly downregulated genes in 2× MIC ciprofloxacin-treated D23580 relative to NT.**

**Table S3. Top 20 significantly upregulated genes in 1× MIC azithromycin D23580 relative to NT.**

**Table S4. Top 20 significantly downregulated genes in 1× MIC azithromycin D23580 relative to NT.**

**Table S5. Top 20 significantly upregulated genes in 50% sucrose fraction of ciprofloxacin-treated D23580 relative to NT.**

**Table S6. 20 top downregulated genes in 50% sucrose fraction of ciprofloxacin-treated D23580 relative to NT.**

**Table S7. Top 20 significantly upregulated genes in ciprofloxacin-treated D23580 60% sucrose fraction relative to ciprofloxacin-treated D23580 50% fraction.**

**Table S8. Top 20 significantly downregulated genes in ciprofloxacin-treated D23580 60% sucrose fraction relative to ciprofloxacin-treated D23580 50% fraction.**

## Supplementary data file legends (available online)

**Supplementary data file S1. RNA-seq differential expression analysis results of *S.* Typhimurium isolates SL1344, D23580, and VNS20081.** Filtered (padj < 0.05) and unfiltered DESeq2 results for each isolate (ciprofloxacin-treated relative to untreated) and the common differentially expressed genes between isolates. Sheet 1 (“Table_of_contents”) provides sheet names and descriptions for data included in each sheet. Data can be found here: https://doi.org/10.17605/OSF.IO/N9CW5

**Supplementary data file S2. RNA-seq differential expression analysis results of *S.* Typhimurium D23580 exposed to 4 parallel conditions.** Filtered (padj < 0.05) and unfiltered DESeq2 results for each condition (0.5x MIC ciprofloxacin, 2× MIC ciprofloxacin, 1 μg/ml mitomycin C, 1× azithromycin) relative to untreated (NT). Sheet 1 (“Table_of_Contents”) provides sheet names and descriptions for data included in each sheet. Data can be found here: https://doi.org/10.17605/OSF.IO/N9CW5

**Supplementary data file S3. RNA-seq differential expression analysis results of *S.* Typhimurium D23580 sucrose gradients.** Filtered (padj < 0.05) and unfiltered DESeq2 results for each measured sucrose concentration (50% or 60%) and condition (ciprofloxacin-treatment or no treatment (NT)). Sheet 1 (“Table_of_contents”) provides sheet names and descriptions for data included in each sheet. Data can be found here: https://doi.org/10.17605/OSF.IO/N9CW5

**Supplementary data file S4. Sample names and corresponding accession numbers for raw sequencing data stored in ENA.** Table of RNA-sequencing and whole genome sequencing (WGS) sample names and accession numbers for access to data submitted to the European Nucleotide Archive (ENA). Data can be found here: https://doi.org/10.17605/OSF.IO/N9CW5

